# Monthly macroalgal surveys reveal a diverse and dynamic community in an urban intertidal zone

**DOI:** 10.1101/2024.09.12.612793

**Authors:** Siobhan Schenk, Varoon P Supratya, Patrick T Martone, Laura W Parfrey

**Affiliations:** Biodiversity Research Center, Department of Botany, University of British Columbia, 6270 University Blvd, Vancouver, British Columbia V6T 1Z4, Canada; Biodiversity Research Center, Department of Zoology, University of British Columbia, 6270 University Blvd, Vancouver, British Columbia V6T 1Z4, Canada; Hakai Institute, PO Box 25039, Campbell River, British Columbia V9W 0B7, Canada

**Keywords:** Macroalgae, Seaweed, Urban Ecology, Long-Term Survey, Biodiversity monitoring, Phenology, Monthly Sampling, British Columbia, Baseline data, Invasive species

## Abstract

Baseline data are critical to measuring how communities shift in response to climate change and anthropogenic activity. Macroalgae are marine foundation species that are vulnerable to climate change, but baseline macroalgal biodiversity data are lacking for many areas of British Columbia, particularly at a high temporal resolution over years. This presents an obstacle for measuring how communities change in response to shifting average conditions as well as after extreme events, such as the 2021 heat dome. To increase baseline macroalgae biodiversity data in British Columbia, we established a monthly transect-based macroalgal survey in 2021 at an urban intertidal site with a low-cost and easily replicable survey protocol, in addition to publishing a previously unpublished 1983-1984 historical dataset of the same area. Our datasets and our survey protocol are freely available. Over 35 months we have recorded 61 taxa of macroalgae, including the canopy forming kelp *Nereocystis luetkeana* and the introduced fucoid *Sargassum muticum*. Surveying throughout the year at regular intervals has revealed large-scale seasonal shifts in macroalgal community composition, the timing of kelp recruitment, and a decrease in abundance of *Fucus distichus* in the upper intertidal zone over multiple years. Our publicly accessible data constitute the most comprehensive survey of intertidal macroalgal diversity in Burrard Inlet, illustrating how simple surveying methods can provide high resolution records of macroalgae biodiversity, particularly in easily accessible urban environments.

## Introduction

Marine macroalgae (seaweeds) are highly diverse and globally distributed foundation species. Macroalgae create habitat for other organisms (Steneck et al. 2002; Teagle et al. 2017), physically and chemically modify their surrounding environment (Pfister et al. 2019), and support higher trophic levels through primary production (Dayton 1985; Mann 1973). Macroalgae are also economically valuable, contributing 500 billion USD per year to the global economy, through ecosystem services, wild harvesting, and mariculture (Eger et al. 2023). Macroalgae are sensitive to changing environmental conditions, including those resulting from anthropogenic activity (Bindoff et al. 2019; Harley et al. 2012; Harley and Paine 2009). Rising average and extreme sea and air temperatures are the most pervasive anthropogenic impacts on the global biosphere (Lima and Wethey 2012; Parmesan and Yohe 2003; Poloczanska et al. 2013). However, pollution (Vergés et al. 2020), sediment runoff (Blain et al. 2021), eutrophication (Ducrotoy 1999), and invasive species (Salvaterra et al. 2013; Casas et al. 2004) are additional major sources of anthropogenic stress. These stressors all interact to affect macroalgal growth, reproduction, and survival (Breeman, 1988; Davison and Pearson, 1996; Harley et al. 2012; Helmuth et al. 2006), which in turn alters local intertidal range limits (Harley et al. 2012; Harley and Paine 2009; Helmuth et al. 2006), and global distribution (Andersen et al. 2013; Johnson et al. 2011; Steneck et al. 2002; Wernberg et al. 2010; Helmuth et al. 2006).

Shifts in macroalgal biodiversity brought on by climate change and other anthropogenic stressors are likely to have disproportionately large impacts on ecosystem function, such as reduced ecological facilitation (Halpern et al. 2007), habitat loss (Lilley and Schiel 2006), and trophic cascades (Rogers-Bennett and Catton, 2019). Quantifying the effects of anthropogenic stress on macroalgae requires documenting baseline diversity and long-term monitoring to track changes in macroalgal communities over time (Bates et al. 2009; Lindstrom et al. 2021). However, macroalgae remain understudied in comparison to other ecologically important groups (Harley et al. 2012; Wernberg et al. 2012), and comprehensive baselines are lacking in many regions (Bringloe and Saunders 2019; Mineur et al. 2015). The lack of macroalgal baseline data was apparent in British Columbia (BC) following the July 2021 heat dome, which subjected exposed intertidal organisms to temperatures of 43°C or higher in BC (White et al. 2023). With the available baseline data for intertidal invertebrates, heat dome mortality rates of over 70% could be estimated for mussels and barnacles (White et al. 2023), yet baseline data to make analogous assessments for local macroalgae were lacking despite anecdotally observed mass die-offs

In BC, macroalgal communities have often been surveyed along remote, relatively exposed coastlines. Notable published macroalgal surveys have occurred on Calvert Island (Whalen et al. 2023), the West coast of Vancouver Island (Starko et al. 2019), Haida Gwaii (Levings and Stewart 2020), and the Gulf islands (Demes et al. 2012; Lindstrom 1973). In comparison, the published literature on macroalgal communities near urbanized regions has been more limited, even though these areas experience comparatively greater levels of anthropogenic stress, including pollution, sedimentation, abiotic extremes, and invasive species (Blain et al. 2021; Ducrotoy 1999; Heery et al. 2018; Todd et al. 2019; Vergés et al. 2020). A previous study in Howe Sound (Wonham et al. 2023), which has experienced extensive industrial activity, combined historical datasets and two sampling events from June to July in 2015. Another study in Burrard Inlet documented the diversity and phenology of intertidal kelps in 1969 and 1970 but did not record other macroalgae (Druehl and Hsiao 1977). A 1983-1984 monthly survey by David J. Garbary recorded intertidal and subtidal macroalgal diversity and seasonality at multiple sites around metropolitan Vancouver, but was unpublished.

Addressing this knowledge gap, we initiated a macroalgal monitoring program in September 2021 at an intertidal site in Burrard Inlet, located in southern British Columbia. The highly urbanized character of Burrard Inlet was intended as a comparison to more isolated regions in BC to understand the impacts of intense anthropogenic stress on macroalgae. Furthermore, the accessibility of Burrard Inlet compared to more isolated study sites made it amenable to high-frequency year-round surveying, facilitating the detection of subtle phenological shifts and seasonal changes in taxonomic composition. Our goal was to establish a comprehensive baseline dataset of: *(i)* Overall diversity of intertidal macroalgae at the site, *(ii)* Presence of any possible introduced, invasive species, such as *Sargassum muticum* (Pawluk 2016; Salvaterra et al. 2013), *Mazzaella japonica* (Pawluk 2016), or *Undaria pinnatifida* (Casas et al. 2004), *(iii)* Spatial patterns of macroalgal diversity across the intertidal gradient, and *(iv)* Intra and interannual shifts in diversity and phenology of conspicuous algae. Alongside our data, we include and publish David J. Garbary’s 1983-1984 historical dataset, to which we compare our transect data to assess *(i)* changes in diversity over time and *(ii)* differences in seasonal trends between historical and current datasets.

## Methods

### Survey site

In September 2021, we started conducting monthly intertidal vertical transect-based macroalgal biodiversity surveys near the Girl in a Wetsuit statue (GW) in Stanley Park, Vancouver, BC, Canada (49.3027, −123.1265; Fig. 1A). GW is located in the First Narrows region of Burrard Inlet, which is heavily urbanized and located near Downtown Vancouver and the Port of Vancouver, Canada’s largest port (Cargo and Terminals 2015). This site is a well-known local hotspot for intertidal biodiversity and has been extensively utilized by numerous research groups and educators from local institutions including the University of British Columbia and Simon Fraser University, as well as various non-academic stakeholders (e.g., Stanley Park Ecological Society). While comparatively undeveloped, Stanley Park has also been subject to extensive shoreline modification, and experiences substantial foot, bicycle, vehicle, and equine traffic. Water flow in Burrard Inlet has been significantly impacted by development (Tsleil-Waututh Nation 2017). The First and Second Narrows experience cooler and more saline surface waters than in the remainder of Burrard Inlet due to turbulent mixing as tidal flows pass through these constrictions (Davidson 1973). This difference in salinity is particularly evident in the spring, when the freshwater outflow from nearby major rivers is greatest, emptying directly into Burrard Inlet (Neilson-Welsh 1999). The side of Stanley Park where GW is located maintains salinities above 20 ppt, while nearby regions drop as low as 10 ppt (Ryan et al. 2019; Druehl and Hsiao 1977). The intertidal zone at GW consists of a near vertical seawall over a gently sloping beach of mixed cobble interspersed with occasional large boulders. The beach extends up to 95 m from the seawall when low tide reaches 0 m chart datum (Fig. 1B).

**Figure 1.**
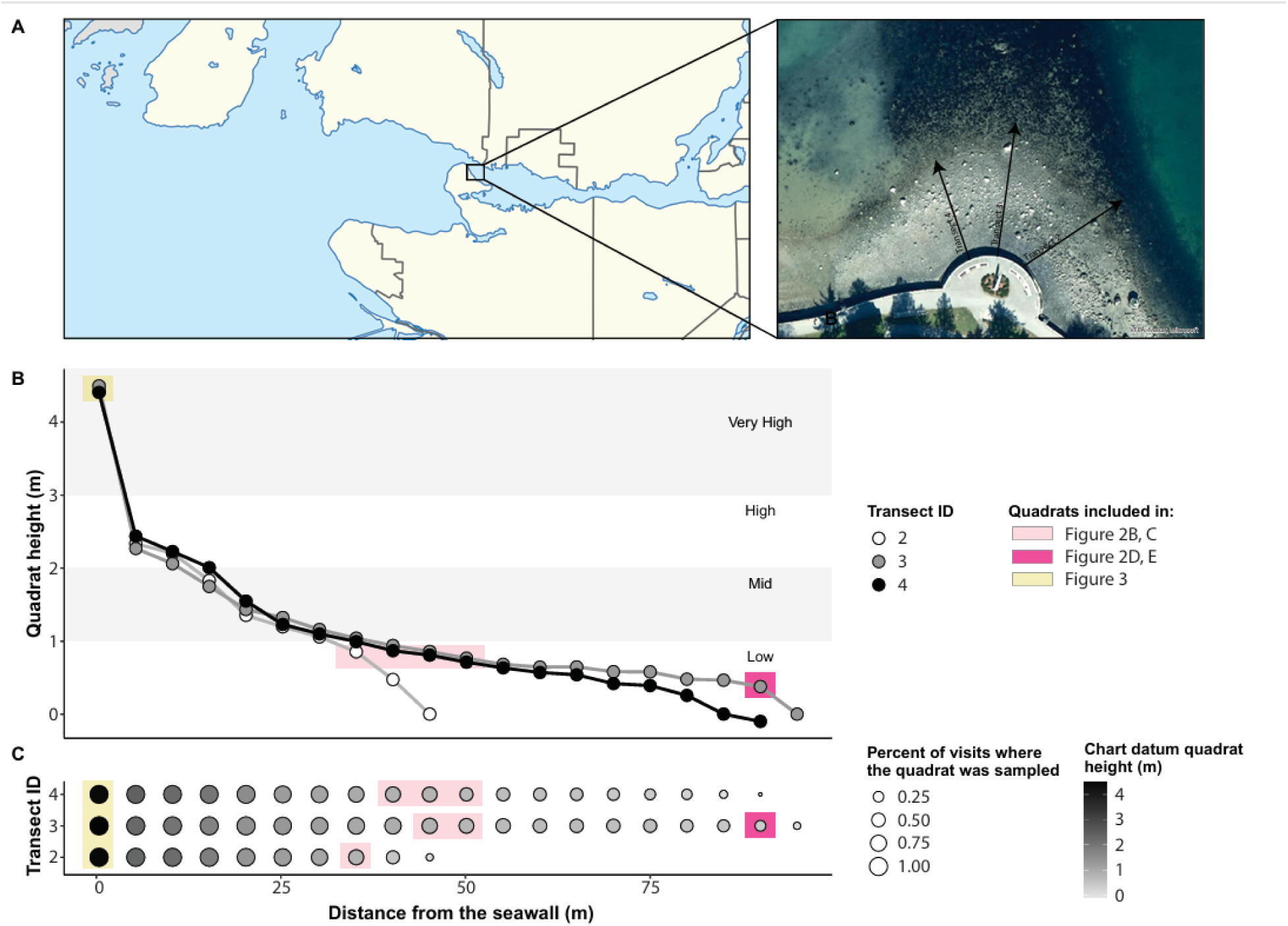
**A)** Map of local area with inset showing overhead photo and position of transects. **B)** The chart datum quadrat height and distance from the seawall along each of the three transects, showing the slope of the site. Rough delineations of intertidal zones based on shore height are indicated **C)** The frequency at which each quadrat was sampled as a percent of total sampling events (size) colored by chart datum quadrat height in metres. Note, the blue boxes in panels B and C indicate the six quadrats between 0.7 and 0.9 m chart datum height. The brown box in panels B and C show the quadrat at 90 m of transect 3.

### Survey design

Our survey methodology was designed to be low cost and easy to replicate, while maximizing the chances of capturing changes in intertidal macroalgal communities over time, particularly distributional and phenological shifts that are expected to occur in response to climate change. The accessibility of GW permits a high-frequency monthly survey design that provides increased phenological resolution, which is critical because phenological shifts tend to be relatively subtle (measured in days to weeks; Fitter and Fitter 2002; Rawal et al. 2015), and therefore require frequent monitoring to detect.

The monthly survey comprised three vertical transects, running perpendicularly to the shore from the seawall towards the water (Fig. 1B). Vertical transects are a well-established survey design for ecological surveys in intertidal habitats and are well suited to capture range shifts along the intertidal gradient across intra- and interannual timescales (Druehl and Green 1982; Miller and Ambrose 2000). The transect lines were replicated monthly using multiple photographically documented spatial reference points. On the seawall, these consisted of specific, conspicuous landmarks that are part of the seawall structure. Lower in the intertidal zone, the spatial reference points were very large boulders that are unlikely to move between survey events. The length of the transects varied between survey events because the accessible intertidal zone depends on tidal height (Fig. 1C). In general, we conduct each survey at the lowest tide of the month, though this has not always been possible.

### Transect data collection protocol

Starting at the 0 m mark (at the seawall), 1 m^2^ square quadrats were positioned every 5 m on the right-hand side of the transect (directions from the perspective of an observer looking down the shore towards the water) until the water’s edge. The quadrats at 0 m were oriented vertically against the seawall, while all other quadrats were laid flat on the ground. After positioning the quadrat, a reference photo was taken and the percent cover of each macroalgal morphospecies attached to substrate within the quadrat was estimated as a metric of abundance. If the estimated percent cover was between 0% and 5%, percent cover was estimated to the nearest percent. If the estimated percent cover was over 5%, percent cover was estimated to the nearest 5%. Therefore, our percent cover values were: 0, 1, 2, 3, 4, 5, 10, 15, …, 100.

Surveyor team size and composition varied by sampling visit and by season. In the winter, when macroalgal diversity and abundance were relatively low and the transect distances relatively short, the two co-first authors were generally able to survey all three transects on their own within roughly two hours. In the summer, particularly from May to July, macroalgal diversity and abundance were relatively high, which increased the time to survey each quadrat. During this time, up to four additional surveyors (the two senior authors on this manuscript and the individuals listed in the acknowledgements) assisted with each survey. To complete the surveys from May to July, a minimum of three surveyors (1 per transect) were required. We provided a custom-made field guide with photos and key morphological characteristics to help less experienced surveyors identify the most common macroalgae at the site.

As in Bates et al. (2009), macroalgal morphospecies were identified in the field when possible. Macroalgae that were not identifiable in the field were brought to the University of British Columbia and identified to the lowest taxonomic level possible using a morphology-based taxonomic key (Gabrielson and Lindstrom 2018). Macroalgal specimens that were still ambiguous, had an uncertain taxonomic status or that required genetic analysis to identify (e.g., crustose coralline algae) were assigned to a lumped morphogroup or identified to genus (see taxonomy notes in the supplemental methods). Genus-level taxonomic assignments retain most of the taxon richness information despite requiring significantly less identification expertise (Bates et al. 2007). A taxon list is available in Table 1. As a reference for future verification of taxonomic identifications, a voucher collection of representative specimens for each macroalgal morphospecies was accessioned into the Beaty Biodiversity Museum Algae Collection (accession list on Borealis). Before uploading the macroalgal list to Borealis, macroalgae that were only recorded once across all quadrats and survey events and not verified in the lab were removed. Currently, this resulted in *Plocamium* sp. being excluded from the published dataset.

**Table 1.**
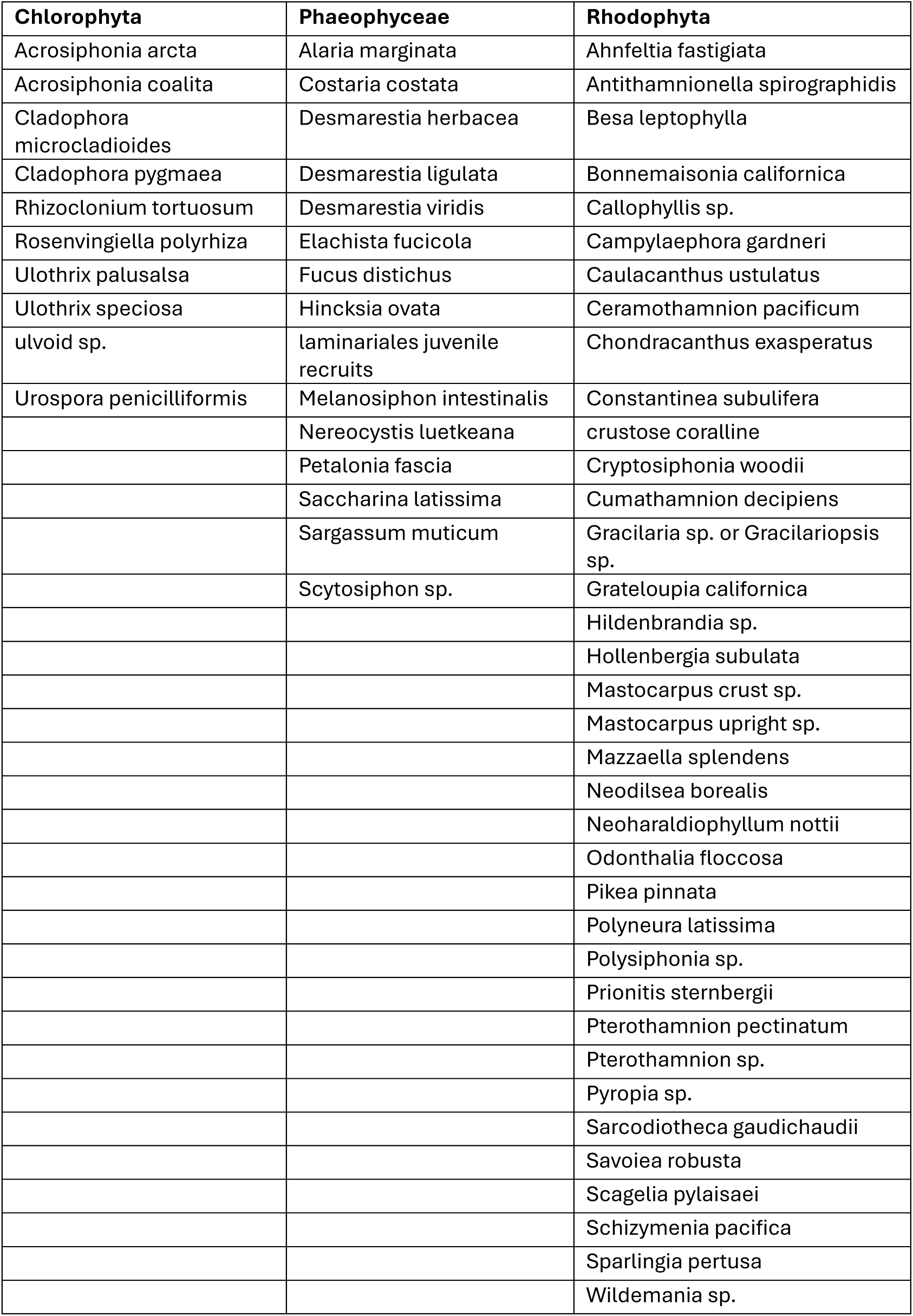
Species list.

Quadrat tidal heights (Fig. 1) were measured on August 1^st^ and 13^th^, 2023 with stadia poles and a bubble level. The quadrat tidal heights were recorded from the lower corner of the quadrat, touching the transect (lower left quadrat corner). Four previously surveyed quadrats (transect 2, 45 m; transect 3, 95 m; transect 4, 85 m and 90 m) were underwater when quadrat tidal heights were measured, and thus inaccessible. The tidal height of these quadrats was extrapolated based on known tide heights from previous surveys when these quadrats were accessible for surveying. Quadrat tidal height measurements are indicated as real or extrapolated in the published dataset.

### Kelp reproductive timing

Starting in April 2022, we opportunistically recorded the reproductive status of the four Laminariales (kelp) species present at GW. Reproductive status was recorded as a binary variable indicating if any thalli with sori were observed anywhere at GW during the survey, as performed in Druehl and Hsiao (1977), providing rough reproductive phenology data for each kelp species. We did not conduct subtidal surveys, so we could not record reproductive status of subtidal kelps.

### Data analysis

Data were analyzed in R 4.4.0 (R Core Team 2022). Measures of central tendency were calculated with the plyr package, version 1.8.9 (Wickham 2011). Plots were generated with ggplot2, version 3.5.0 (Wickham 2016) and formatted with ggpubr, version 0.6.0 (Kassambara 2022).

### Publication and analysis of 1983-1984 algal survey from nearby sites

A dataset from 1983-1984 collected by David J. Garbary has been used by many researchers from BC as baseline data for the region, but these data were not publicly accessible online. These data are monthly (or less frequent depending on the site) presence/absence surveys of multiple sites around Metro Vancouver including both macro- and microalgae. These data include dive surveys.

We updated the historical names in the dataset to reflect contemporary taxonomy, while also retaining the original name assignment. The taxonomy of the dataset was updated using AlgaeBase (Guiry and Guiry 2024) as a primary reference and Gabrielson and Lindstrom (2018) as a secondary reference. Where there were discrepancies between the two references, the reasoning provided in the footnotes of Gabrielson and Lindstrom (2018) was usually sufficient to resolve the discrepancy. In cases where this was not possible, the taxonomy in AlgaeBase (Guiry and Guiry 2024) was followed.

We analyzed a subset of the historical dataset for comparisons with our contemporary baseline data. We removed microalgae and endophytes, as our survey did not capture these. We also restricted our comparisons to four sites around Stanley Park (Third Beach, Siwash Rock, Brockton point, Coal Harbour), which were closest to our contemporary survey site. After these data cleaning steps, we compared our macroalgal list to this 1983-1984 list. We analyzed the 1983-1984 taxon richness following the same procedure as for our transect data.

## Results

### Macroalgal biodiversity was high and seasonally dynamic in urban environments

In total, we identified 61 macroalgal taxa over the entire survey duration (Table 1) These included 15 brown algae (Phaeophyceae), 10 green algae (Chlorophyta), and 36 red algae (Rhodophyta). Across all years, we recorded the lowest mean macroalgal richness (± se) in September (6.0 ± 1.7), closely followed by October (6.7 ± 1.9). We recorded the highest mean macroalgal richness in May (34.0 ± 0.6), followed by June (30.3 ± 0.3). Red algae always had the greatest richness, followed by the brown, then green algae (Fig. 2A). Taxon richness and seasonal trends were not significantly different between transects when controlling for tidal height. Our observed taxon richness was lower than in the 1983-1984 historical dataset; we re-encountered 12 out of 20 brown, 6 out of 12 green, and 30 out of 72 red algal taxa present in the historical dataset. While the vast majority of taxa observed in our contemporary data were present in the historical dataset, we documented 2 brown, 3 green, and 5 red algal taxa that had not been previously recorded. Seasonal changes in diversity were similar between historical and contemporary datasets.

**Figure 2.**
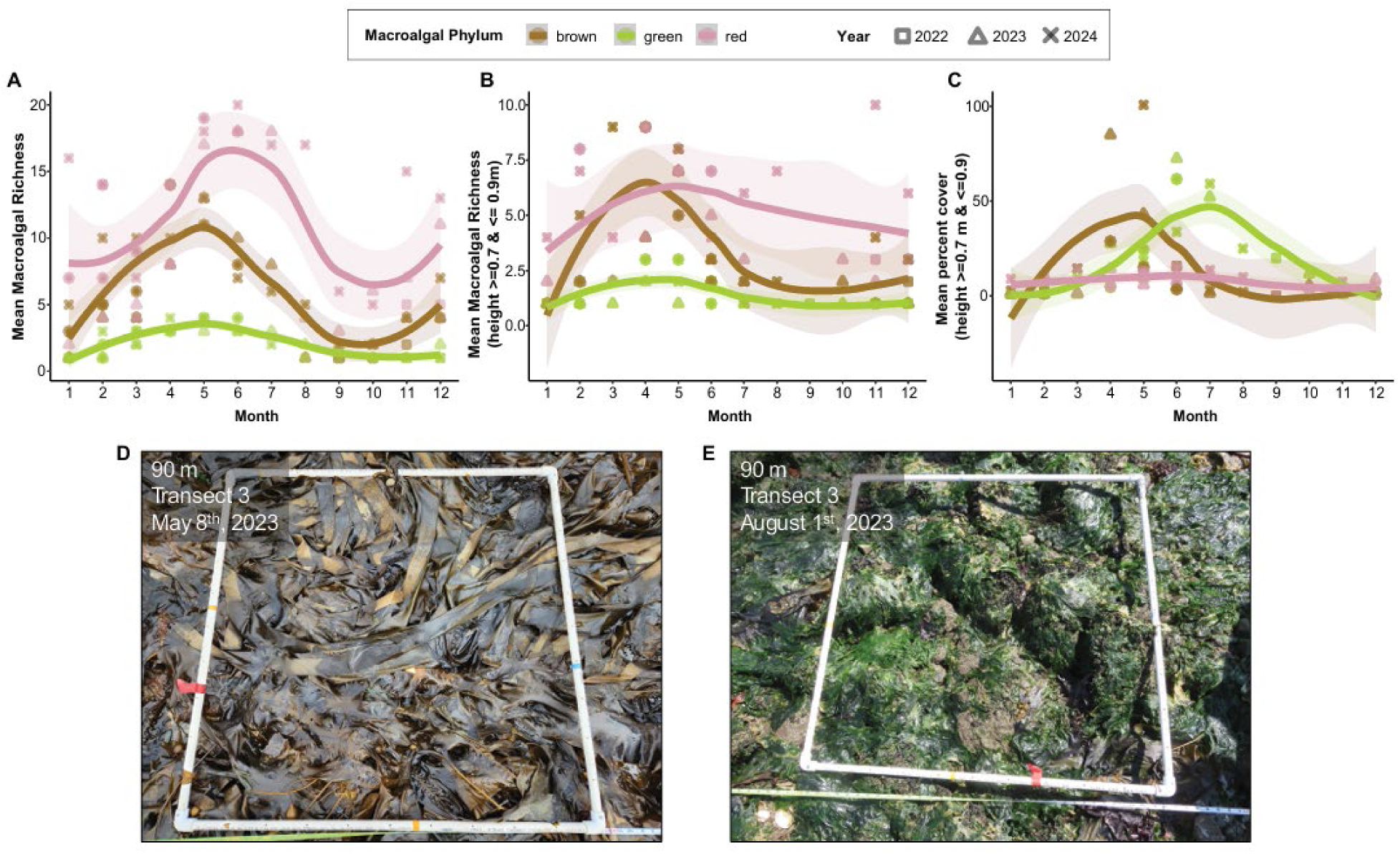
Line graphs showing the mean richness of each macroalgal group by across (A) the entire site or (B) between 0.7 and 0.9 m chart datum. (C) The mean macroalgal percent cover between 0.7 to 0.9 m chart datum. Transparent ribbons represent the standard error around the mean. Representative images from (D) May 2023 and (E) August 2023 of the change from high brown algal cover, dominated by kelps, to high green algal cover, dominated by ulvoid algae. Both images are from transect number 3, 90 m from the seawall (0 m chart datum height).

The taxon richness observed in our overall dataset appears to be negatively correlated with tidal height in the intertidal zone. We were able to access lower tide heights in the spring and early summer compared to fall and winter, potentially biasing the results towards higher observed taxon richness in spring and summer (Fig. 2A). To control for the effect of tide height on taxonomic richness, we confined our calculations to the lowest six quadrats that were regularly sampled year-round, which were between 0.7 and 0.9 m chart datum (Fig. 1C,2B). Between 0.7 and 0.9 m chart datum, seasonal changes in taxon richness were less pronounced for all major macroalgal groups than in the overall dataset, particularly for red and green algae. However, all major macroalgal groups still exhibited a springtime peak in richness, particularly for brown algae (Fig. 2B), matching the kelp recruitment period. In contrast to the seasonal dynamics we observed intertidally, the 1983-1984 Garbary data, which included subtidal surveys unaffected by sampling tidal height, displayed minimal seasonal variation in taxonomic richness.

### Macroalgal cover varied seasonally

The percent cover of macroalgae also exhibited substantial seasonal variation (Fig. 2C). Notably, we observed a striking annual shift from a brown algae dominated intertidal zone in the spring and early summer, to a green algae dominated intertidal zone in midsummer to fall, while overall macroalgal cover was low in winter (Fig. 2C). This seasonal cycling was the result of a springtime period of kelp recruitment and growth. Starting in February, young kelp recruits (particularly *Saccharina latissima, Alaria marginata,* and *N. luetkeana*) could be found throughout the entire intertidal zone as high as the seawall on all substrate types (boulders, cobble and shell hash). By April to May, kelps collectively reached 50 to 100% cover in the lower intertidal zone (Fig. 2C,D). Kelps then progressively died back to the subtidal following warm summer low tides. The kelp dieback was followed by an increased abundance of heat- and desiccation tolerant green ulvoid algae (Fig. 2C,E). Ulvoid algae persisted in high abundance into the later summer and early fall, and thereafter declined gradually into the winter. Despite red algae being much more species-rich than the green and brown algae combined (Fig. 2B), average red algal cover was low and varied little over the year even after survey height in the intertidal zone was controlled for (Fig. 2C).

Individual macroalgal taxa also showed strong seasonal patterns in abundance, influencing broader macroalgal richness trends. For example, the brown algae *Melanosiphon intestinalis* and *Petalonia fascia* were recorded throughout the year but had a significantly higher cover (up to 10 times higher) from February to May. *Scytosiphon* sp., *Acrosiphonia arcta*, and *Acrosiphonia coalita* were found across a large range of the intertidal zone at GW, but only between March and June.

### Decreasing macroalgal cover in the upper intertidal zone

We recorded a large decline in macroalgal cover in the upper intertidal zone starting around in November 2022 (Fig. 1C, 3A). Much of this decline was due to the decline of *Fucus distichus* (Fig. 3B,C,D,E). *F. distichus* cover did not decline lower on the shore (Fig. S1), showing that the decline in percent cover was confined to the seawall, rather than a site-wide phenomenon. Other macroalgae in the high intertidal zone, such as *Mastocarpus* sp. gametophytes (upright stage), showed no significant change in cover during the same time (Fig. 3B). Fleshy crustose algae (*Hildenbrandia* sp. and sporophytes of M*astocarpus* sp.) increased in abundance on the seawall during the *F. distichus* decline and generally throughout the duration of the survey (Fig. 3B). In fact, fleshy crustose algae were absent from the seawall until March 2022, though they were recorded below 0.6 m chart datum since December 2021.

**Figure 3.**
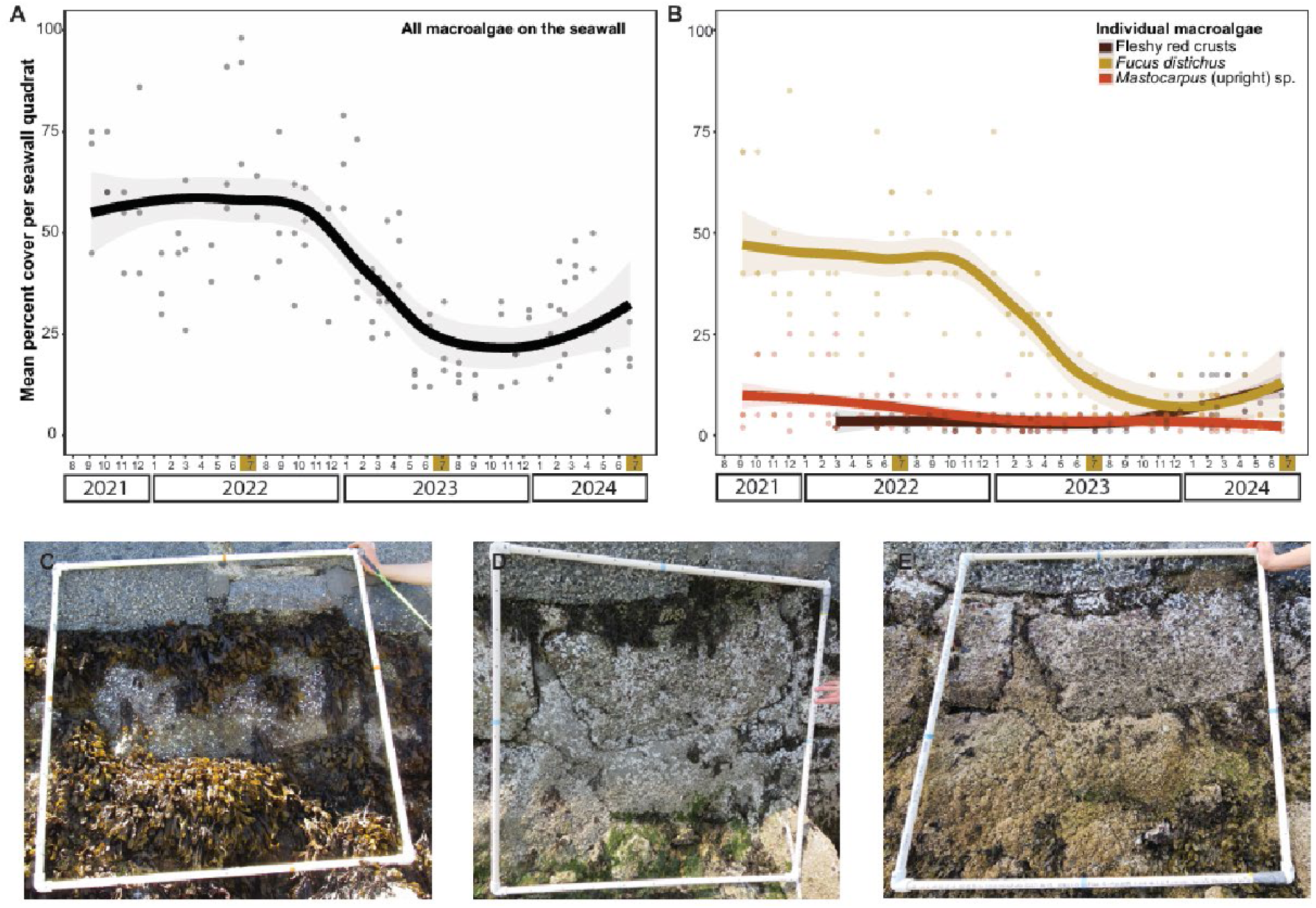
Plot of the percent macroalgal cover per quadrat on the seawall of (A) all macroalgae and (B) selected macroalgal species. Points show the individual transect values and the lines represent the mean of all three transects. Ribbons show the standard error around the mean. (C,D,E) Photos from the seawall quadrat from transect 3 in July C) 2022, D) 2023, and E) 2024. Brown squares on the x-axis of panels A and B correspond to the dates shown in photos C to E.

### Kelp reproductive timing data

We observed all four kelp species present intertidally at GW having sori or sporophylls at least once throughout our monitoring program (Fig. 4). Reproductive *S. latissima* was found in all months except January, March, and June. *N. luetkeana* showed a clear trend in reproductive timing, with sori appearing between April and September. Reproductive *Costaria costata* and *A. marginata* were rarely observed reproducing intertidally, with three and one observed instance(s) respectively, all in 2024 (Fig. 4).

**Figure 4.**
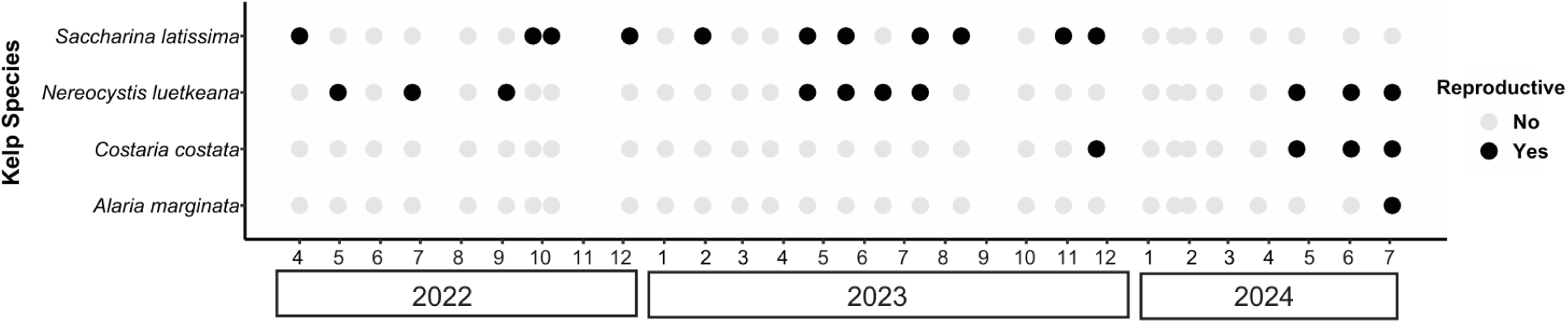
Plot showing if any reproductive kelp by species were observed by sampling visit.

## Discussion

A lack of baseline data is a significant obstacle for quantifying changes in macroalgal communities in response to climate change and other anthropogenic stressors (Bates et al. 2009; Lindstrom et al. 2021). Our macroalgal monitoring program documented the diverse and dynamic intertidal macroalgal community at an urban intertidal site in BC following an extreme weather event (2021 heat dome), not only contributing to an improved understanding of the ecology of Burrard Inlet, but also serving as a baseline against which to track the future impact of ongoing anthropogenic activity (Bindoff et al. 2019; Poloczanska et al. 2013; Wonham et al. 2023). Between September 2021 and July 2024, we recorded a total of 61 taxa of macroalgae and captured intra- and interannual trends in community composition, including seasonal shifts in dominant macroalgal taxa and a progressive multi-year decline of the foundational *F. distichus* in the upper intertidal zone. The publicly available protocols and easy to update, public dataset we have established provide an opportunity to continue tracking macroalgal communities, helping to improve our understanding of the natural variation and long-term directional trends in biodiversity between years. As of the publication date of this manuscript, the surveys are ongoing and the online data are updated accordingly.

The interannual changes we observed in *F. distichus* abundance in the high intertidal zone highlight the potential for even limited time-series data to capture change. Starting in November 2022, *F. distichus* cover on the seawall underwent a marked decline, and has not recovered as of July 2024. Declines of *F. distichus* have been documented in response to physical disturbance of the shoreline (Klinger and Fukuyama 2011) and marine heatwaves (Suryan et al. 2021; Weitzman et al. 2021; Whalen et al. 2023). However, the decline observed at GW did not appear to be closely associated with any specific trigger, beginning more than a year after the 2021 heat dome. Populations of *Fucus* spp. have been documented to undergo natural, cyclical changes in abundance over multi-year time periods (Klinger and Fukuyama 2011; Little et al. 2017), possibly due to biotic interactions (Little et al. 2017). It is possible that the changes in *F. distichus* abundance we observed represent a portion of a larger, cyclical pattern of variation. However, a longer period of surveying would likely be required to clearly distinguish a decline due to a discrete trigger from cyclical oscillations, as the latter may play out over decadal timeframes (Klinger and Fukuyama 2011; Little et al. 2017).

Compared to our contemporary baseline data, diversity was substantially higher in the historical 1983-1984 dataset for red, green, and brown algae. However, the historical dataset also included subtidal taxa observed during dive surveys covering a variety of substratum types, while we only recorded attached algae in the predominantly cobbly intertidal zone. This likely accounts for much of the greater taxon diversity observed in the historical data. For instance, both co-first authors have observed *Opuntiella californica*, *Sarcodiotheca furcata*, and *Smithora naiadum* (present in the historical data) as drift at GW, but have never observed it attached intertidally. Most other algae present in only one or the other dataset were filamentous and/or inconspicuous, and therefore may have simply been overlooked by historical or contemporary surveyors. Overall macroalgal taxonomic richness at GW does not appear to have meaningfully changed over the past 40 years.

Beyond documenting the overall presence and spatial distribution of macroalgal taxa at GW, our high sampling frequency has also allowed us to capture seasonal dynamics of macroalgal diversity, providing a temporal baseline against which to compare any future shifts in seasonal phenology. One particularly striking seasonal dynamic observed at GW was the mass recruitment of young kelp sporophytes in spring, followed by their subsequent dieback and replacement with green ulvoid algae throughout the summer. The large range of tidal height and substrates on which kelp sporophytes could be found indicate that spore settlement and sporophyte recruitment may be relatively nonselective and not tied to specific substrates. While other studies have suggested that kelp recruitment may be mediated by coralline algae (Barner et al. 2016; Milligan and DeWreede 2000; Okamoto et al. 2013; Twist et al. 2024), they were unlikely to have played a significant positive or negative role as a settlement cue at GW due to the extreme rarity of crustose corallines at GW and absence of geniculate corallines, as well as the fact that kelp recruitment occurred in much higher regions of the intertidal zone than there were coralline crusts. The springtime peak in recruitment observed at GW is consistent with the known population biology of all four kelp species involved: *S. latissima* (Andersen 2013; Nielsen et al. 2014; Raymond 2020; Ulaski et al. 2020), *C. costata* (Maxell and Miller 1996), *A. marginata* (McConnico and Foster 2005; Raymond 2020), *N. luetkeana* (Connell 1961; Dobkowski et al. 2019; Maxell and Miller 1996; Ulaski et al. 2020). However, overall kelp cover in this intertidal study peaked in May and declined thereafter, while previous subtidal studies tended to document kelps persisting into the late summer and autumn (Helmuth et al. 2006). The early dieback of intertidal kelps at GW corresponds to a seasonal shift from cool temperatures and nighttime low-tides (October to March) to warmer temperatures and daytime low tides (April to September) common across the Northeast Pacific that exposed the intertidal zone to increased thermal and desiccation stress (Druehl and Hsiao 1977). The concurrent increase of ulvoid green algae to replace the kelps is consistent with their greater ability to tolerate these abiotic stressors (Thom and Albright 1990; Gao et al. 2017; Nelson, Nelson, and Tjoelker 2003; Gao et al. 2011). Summer blooms of ephemeral ulvoid algae are common in the Salish Sea, where they may overgrow other marine macrophytes and alter biodiversity (Nelson et al. 2008, 2003; Van Alstyne 2016). In Norway, *S. latissima* beds degraded by ocean warming have been shown to seasonally alternate with low-growing turf algae in a similar fashion to the pattern observed at GW, with increased kelp abundance in the spring and turf dominance in the summer (Christie et al. 2019; Moy and Christie 2012).

Intensifying thermal and desiccation stress due to ongoing climate change could accelerate the timing and/or reduce extent of intertidal kelp recruitment, which may be accompanied by an increase in the overall predominance of ulvoid algae at GW. In fact, comparisons with previously published historical data available in the local area suggest that kelp zonation and phenology may already have shifted over the last several decades (Druehl and Hsiao 1977). For instance, while historical data shows that *C. costata* used to recruit as high as 1.3 m chart datum (Druehl and Hsiao 1977), we never encountered *C. costata* above 0.8 m chart datum. We also have observed only a single occurrence of intertidal *A. marginata* bearing sporophylls in three years of monthly surveying, whereas the historical data records reproductive intertidal *A. marginata* from April to September (Druehl and Hsiao 1977).

*Sargassum muticum* was the only invasive macroalga we detected at GW. This species may have been introduced to BC as early as 1902 (Pawluk, 2016), and is known to be established and common throughout the province. At GW, *S. muticum* cover was generally low, at least in the intertidal zone. However, climate change may increase its competitive advantage over native species due to its wide thermal tolerance range compared to many native seaweeds (Atkinson et al. 2020; de la Hoz et al. 2019; Olabarria et al. 2013). We have not detected other macroalgal invasives at GW, such as the red alga *M. japonica* or the kelp *U. pinnatifida*. However, *M. japonica* has already been documented in the Strait of Georgia (Pawluk 2016; Saunders and Millar 2014), and environmental conditions in BC are suitable for the establishment of *U. pinnatifida* (Silva et al. 2002; Zabin et al. 2009), which is established in Northern California (Félix-Loaiza et al. 2022). Regular biodiversity surveys near potential sites of introduction (e.g., major ports, such as the Port of Vancouver) could assist in the early detection of future species invasions.

## Acknowledgements

Dr. B. Clarkston for her invaluable help to identify macroalgae and suggestions on how to improve the study and data usability.

Volunteers who donated their time to help us collect these data. In particular: A. Choinski, A. Jackman, A. Palacios, A. Simon, B. Clarkston, C. Wardrop, E. Evans, E. Jourdain, E. Menchions, E. Kohn, E. Porcher, G. Ainsworth-Cruickshank, MJ. Herrin, N. Salland, P. Lund, P. Schenk, R. Burns, R. Perovich, R. Ogushi, V. Billy, V. Pornsinsiriruk.

Colleagues who helped edit the latest draft of the manuscript: E. Jourdain, R. Burns, R. d’Entremont.

We also thank D.J. Garbary for permitting us to publish his historical dataset.

## Competing interests

The authors declare there are no competing interests.

## Author contributions

Conceptualization: SS, VPS

Data curation: SS

Formal analysis: SS

Funding acquisition: LWP, PTM

Investigation: VPS, SS, LWP

Methodology: SS, VPS, LWP, PTM

Project administration: SS, VPS, LWP

Software: SS

Resources: LWP, PTM

Supervision: LWP, PTM

Visualization: SS, VPS

Writing – original draft: VPS, SS

Writing – review & editing: VPS, SS, LWP, PTM

## Funding

Natural Sciences and Engineering Research Council Discovery Grant (RGPIN-2021-03160) and Canada Research Chair tier 2 (CRC-2019-00252) to LWP. Natural Sciences and Engineering Research Council Discovery grant to PTM (RGPIN-2019-06240). Natural Sciences and Engineering Research Council (NSERC) Postgraduate Scholarship – Doctoral (PGS-D) to VPS (#6564). Botany Graduate Excellence Award #6372, James Robert Thompson Fellowship, Kruger Graduate Fellowship, Ocean Leaders Fellowship (NSERC), and British Columbia Graduate Scholarship to SS.

## Data availability

*Current study:* Detailed sampling protocols, sampling logs, and all data generated during this study are available in the Borealis Dataverse (https://doi.org/10.5683/SP3/IKGB6E), including an accession list of herbarium vouchers of representative macroalgal morphospecies. The dataset may be updated with more sampling timepoints if more sampling occurs past the publication date of this article. The physical herbarium vouchers are deposited at the Beaty Biodiversity Museum (University of British Columbia, Vancouver).

*Garbary 1983-1984 survey:* The data and an overview of survey methods are available on the Borealis Dataverse https://doi.org/10.5683/SP3/3X3L7O.

---

**Figure S1.**
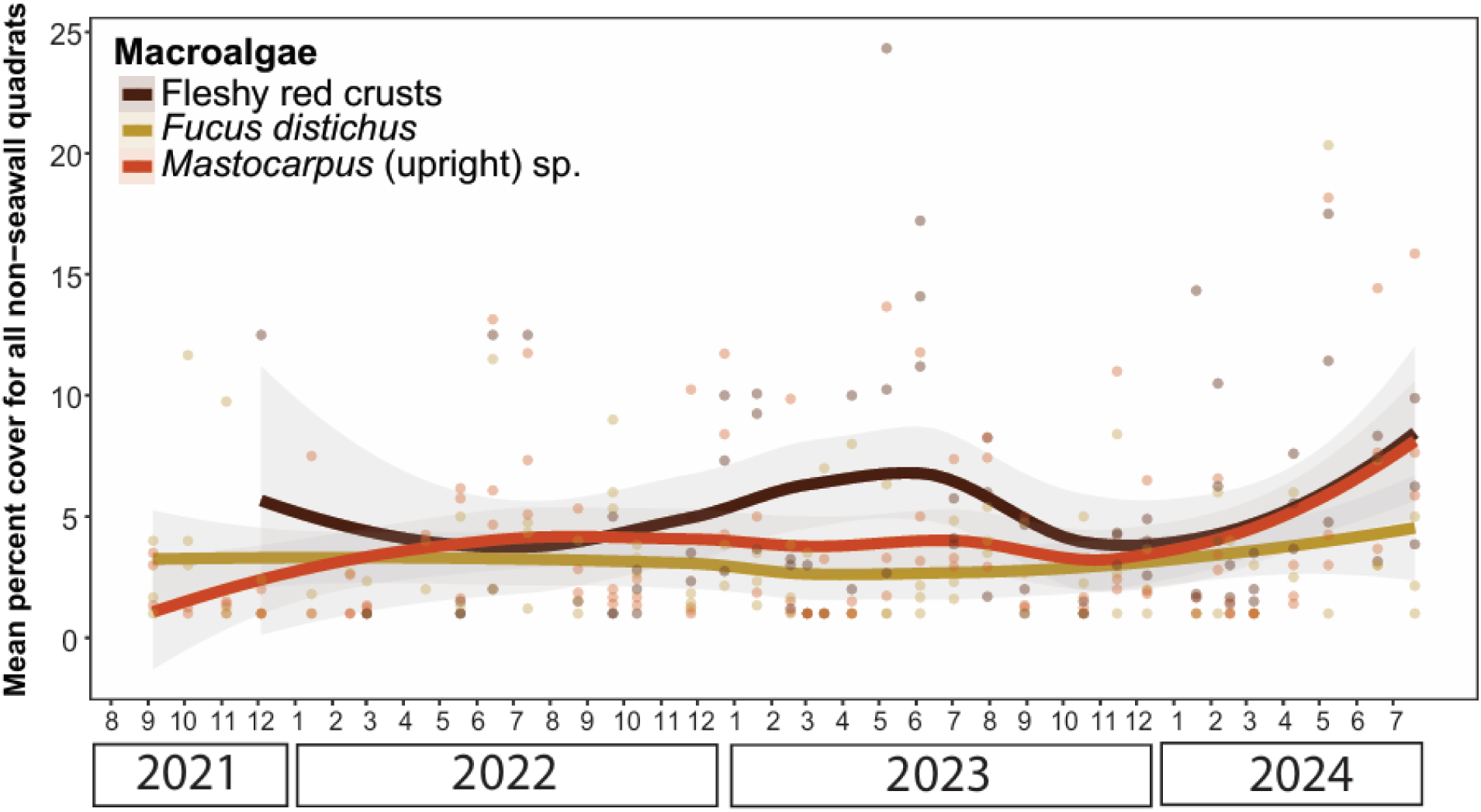
Plot of the percent macroalgal cover per quadrat across all quadrats sampled per sampling visit excluding seawall quadrats (Figure 2B). Points show the individual transect values and the lines represent the mean of all three transects. Ribbons show the standard error around the mean.

## Notes

https://doi.org/10.5683/SP3/IKGB6E

### Competing Interest Statement

The authors have declared no competing interest.

### Summary of Updates

Revisions after peer-review for the journal of Botany

## References

Andersen, G.S. 2013. Patterns of *Saccharina latissima* recruitment. PLoS One. 8(12): e81092. doi:10.1371/journal.pone.0081092.

Andersen, G.S., Pedersen, M.F., Nielsen, S.L. 2013. Temperature acclimation and heat tolerance of photosynthesis in Norwegian *Saccharina latissima* (Laminariales, Phaeophyceae). Phyc. 49: 689–700. doi:10.1111/jpy.12077.

Atkinson, J., King, N.G., Wilmes, S.B., Moore, P.J., 2020. Summer and winter marine heatwaves favor an invasive over native seaweeds. J. Phyc. 56: 1591–1600. doi:10.1111/jpy.13051.

Barner, A.K., Hacker, S.D., Menge, B.A., Nielsen, K.J., 2016. The complex net effect of reciprocal interactions and recruitment facilitation maintains an intertidal kelp community. J Ecol 104: 33–43. doi:10.1111/1365-2745.12495.

Bates, C.R., Saunders, G.W., Chopin, T., 2009. Historical versus contemporary measures of seaweed biodiversity in the Bay of Fundy. Botany 87: 1066–1076. doi:10.1139/B09-067.

Bates, C.R., Scott, G., Tobin, M., Thompson, R., 2007. Weighing the costs and benefits of reduced sampling resolution in biomonitoring studies: Perspectives from the temperate rocky intertidal. Biological Conservation, Forests in the Balance: Linking Tradition and Technology in Lanscape Mosaics. 137: 617–625. doi:10.1016/j.biocon.2007.03.019.

Bindoff, N.L., Cheung, W.W.L., Kairo, J.G., Arístegui, J., Guinder, V.A., Hallberg, R., Hilmi, N., Jiao, M.S. Karim, L. Levin, S. O’Donoghue, S.R. Purca Cuicapusa, B. Rinkevich, T. Suga, A. Tagliabue, and P. Williamson. 2019. Changing ocean, marine ecosystems, and dependent communities. In: IPCC special report on the ocean and cryosphere in a changing climate [H.-O. Pörtner, D.C. Roberts, V. Masson-Delmotte, P. Zhai, M. Tignor, E. Poloczanska, K. Mintenbeck, A. Alegría, M. Nicolai, A. Okem, J. Petzold, B. Rama, N.M. Weyer (eds.)]. Cambridge University Press, Cambridge, UK and New York, NY, USA, pp. 447–587. doi:10.1017/9781009157964.007.

Blain, C.O., Hansen, S.C., Shears, N.T. 2021. Coastal darkening substantially limits the contribution of kelp to coastal carbon cycles. Change Biol. 27: 5547–5563. doi:10.1111/gcb.15837.

Breeman, A.M. 1988. Relative importance of temperature and other factors in determining geographic boundaries of seaweeds: Experimental and phenological evidence. Helgolander Meeresunters. 42: 199–241. doi:10.1007/BF02366043.

Bringloe, T.T., Saunders, G.W. 2019. DNA barcoding of the marine macroalgae from Nome, Alaska (Northern Bering Sea) reveals many trans-Arctic species. Polar Biol. 42: 851–864. doi:10.1007/s00300-019-02478-4

Cargo and terminals. 2015. Port of Vancouver. URL https://www.portvancouver.com/cargo-terminals/ (accessed 1.19.24).

Casas, G., Scrosati, R., Luz Piriz, M. 2004. The invasive kelp *Undaria Pinnatifida* (Phaeophyceae, Laminariales) reduces native seaweed diversity in Nuevo Gulf (Patagonia, Argentina). Biol. Invasions. 6: 411–416. doi:10.1023/B:BINV.0000041555.29305.41.

Christie, H., Andersen, G.S., Bekkby, T., Fagerli, C.W., Gitmark, J.K., Gundersen, H., Rinde, E. 2019. Shifts between sugar kelp and turf algae in Norway: Regime shifts or fluctuations between different opportunistic seaweed species? Front. Mar. Sci. 6: doi:10.3389/fmars.2019.00072.

Connell, J.H. 1961. The influence of interspecific competition and other factors on the distribution of the barnacle *Chthamalus stellatus*. Ecology. 42: 710–723. doi:10.2307/1933500.

Davidson, L.W. 1973. On the physical oceanograhy of Burrard Inlet and Indian Arm, British Columbia. University of British Columbia.

Davison, I.R., Pearson, G.A. 1996. Stress tolerance in intertidal seaweeds. J. Phyc. 32: 197–211. doi:10.1111/j.0022-3646.1996.00197.x.

Dayton, P.K. 1985. Ecology of kelp communities. Annu. Rev. of Ecol. Syst. 16: 215–245. doi:10.1146/annurev.es.16.110185.001243.

de la Hoz, C.F., Ramos, E., Puente, A., Juanes, J.A. 2019. Climate change induced range shifts in seaweeds distributions in Europe. Res. 148: 1–11. doi:10.1016/j.marenvres.2019.04.012

Demes, K.W., Kordas, R.L., Jorve, J.P. 2012. Ferry wakes increase seaweed richness and abundance in a sheltered rocky intertidal habitat. Hydrobiologia. 693: 1–11. doi:10.1007/s10750-012-1082-4.

Dobkowski, K.A., Flanagan, K.D., Nordstrom, J.R. 2019. Factors influencing recruitment and appearance of bull kelp, *Nereocystis luetkeana* (phylum Ochrophyta). J Phycol. 55: 236–244. doi:10.1111/jpy.12814.

Druehl, L., Green, J. 1982. Vertical distribution of intertidal seaweeds as related to patterns of submersion and emersion. Mar. Ecol. Prog. Ser. 9: 163–170. doi:10.3354/meps009163.

Druehl, L.D., Hsiao, S.I.C. 1977. Intertidal kelp response to seasonal environmental changes in a British Columbia Inlet. J. Fish. Res. Bd. Can. 34: 1207–1211. doi:10.1139/f77-176.

Ducrotoy, J. 1999. Indications of change in the marine flora of the North Sea in the 1990s. Mar. Pollut. Bull. 38: 646–654. doi:10.1016/S0025-326X(98)00195-7.

Eger, A.M., Marzinelli, E.M., Beas-Luna, R., Blain, C.O., Blamey, L.K., Byrnes, J.E.K., Carnell, P.E., Choi, C.G., Hessing-Lewis, M., Kim, K.Y., Kumagai, N.H., Lorda, J., Moore, P., Nakamura, Y., Pérez-Matus, A., Pontier, O., Smale, D., Steinberg, P.D., Vergés, A. 2023. The value of ecosystem services in global marine kelp forests. Nat Commun. 14: 1894. doi:10.1038/s41467-023-37385-0.

Félix-Loaiza, A.C., Rodríguez-Bravo, L.M., Beas-Luna, R., Lorda, J., Cruz-González, E. de L., Malpica-Cruz, L., 2022. Marine heatwaves facilitate invasive algae takeover as foundational kelp. Bot. Mar. 65: 315–319. doi:10.1515/bot-2022-0037.

Fitter, A.H., Fitter, R.S.R. 2002. Rapid changes in flowering time in British plants. Science. 296: 1689–1691. doi:10.1126/science.1071617.

Gabrielson, P.W., Lindstrom, S.C. 2018. Keys to the seaweeds and seagrasses of Southeast Alaska, British Columbia, Washington, and Oregon. Sandra C Lindstrom, Department of Botany, University of British Columbia. Ed 2. ISBN: 9780976381730.

Gao, G., Clare, A.S., Rose, C., Caldwell, G.S. 2017. Eutrophication and warming-driven green tides (*Ulva rigida*) are predicted to increase under future climate change scenarios. Mar. Poll. Bull. 114: 439–447. doi:10.1016/j.marpolbul.2016.10.003.

Gao, S., Shen, S., Wang, G., Niu, J., Lin, A., Pan, G. 2011. PSI-driven cyclic electron flow allows intertidal macro-algae *Ulva* sp. (Chlorophyta) to survive in desiccated conditions. Plant and Cell Physiology. 52: 885–893. doi:10.1093/pcp/pcr038.

Guiry, M.D., Guiry, G.M. 2024. AlgaeBase. World-wide electronic publication. Accessed 2024-11-27. URL: https://www.algaebase.org/.

Halpern, B.S., Silliman, B.R., Olden, J.D., Bruno, J.P., Bertness, M.D. 2007. Incorporating positive interactions in aquatic restoration and conservation. Front. Ecol. Environ. 5: 153–160. doi:10.1890/1540-9295(2007)5[153:IPIIAR]2.0.CO;2.

Harley, C.D.G., Anderson, K.M., Demes, K.W., Jorve, J.P., Kordas, R.L., Coyle, T.A., Graham, M.H. 2012. Effects of climate change on global seaweed communities. J. Phyc. 48: 1064–1078. doi:10.1111/j.1529-8817.2012.01224.x.

Harley, C.D.G., Paine, R.T. 2009. Contingencies and compounded rare perturbations dictate sudden distributional shifts during periods of gradual climate change. Proc. Natl. Acad. Sci. U.S.A. 106: 11172–11176. doi:10.1073/pnas.0904946106.

Heery, E.C., Hoeksema, B.W., Browne, N.K., Reimer, J.D., Ang, P.O., Huang, D., Friess, D.A., Chou, L.M., Loke, L.H.L., Saksena-Taylor, P., Alsagoff, N., Yeemin, T., Sutthacheep, M., Vo, S.T., Bos, A.R., Gumanao, G.S., Syed Hussein, M.A., Waheed, Z., Lane, D.J.W., Johan, O., Kunzmann, A., Jompa, J., Suharsono, Taira, D., Bauman, A.G., Todd, P.A. 2018. Urban coral reefs: Degradation and resilience of hard coral assemblages in coastal cities of East and Southeast Asia. Mar. Poll. Bull. 135: 654–681. doi:10.1016/j.marpolbul.2018.07.041.

Helmuth, B., Broitman, B.R., Blanchette, C.A., Gilman, S., Halpin, P., Harley, C.D.G., O’Donnell, M.J., Hofmann, G.E., Menge, B., Strickland, D. 2006. Mosaic patterns of thermal stress in the rocky intertidal zone: Implications for climate change. Ecol. Monogr. 76: 461–479. doi:10.1890/0012-9615(2006)076[0461:MPOTSI]2.0.CO;2.

Johnson, C.R., Banks, S.C., Barrett, N.S., Cazassus, F., Dunstan, P.K., Edgar, G.J., Frusher, S.D., Gardner, C., Haddon, M., Helidoniotis, F., Hill, K.L., Holbrook, N.J., Hosie, G.W., Last, P.R., Ling, S.D., Melbourne-Thomas, J., Miller, K., Pecl, G.T., Richardson, A.J., Ridgway, K.R., Rintoul, S.R., Ritz, D.A., Ross, D.J., Sanderson, J.C., Shepherd, S.A., Slotwinski, A., Swadling, K.M., Taw, N. 2011. Climate change cascades: Shifts in oceanography, species’ ranges and subtidal marine community dynamics in eastern Tasmania. J. Exp. Mar. Bio. Ecol. 400: 17–32. doi:10.1016/j.jembe.2011.02.032.

Kassambara, A., 2022. ggpubr: “ggplot2” Based Publication Ready Plots.

Klinger, T., Fukuyama, A.K. 2011. Decadal-scale dynamics and response to pulse disturbance in the intertidal rockweed *Fucus distichus* (Phaeophyceae). Mar. Ecol. 32: 313–319. doi:10.1111/j.1439-0485.2011.00481.x.

Levings, C.D., Stewart, H.L. 2020. Research priorities for nearshore algae in coastal British Columbia workshop and gap analysis – final report. Fisheries and Oceans Canada.

Lilley, S.A., Schiel, D.R. 2006. Community effects following the deletion of a habitat-forming alga from rocky marine shores. Oecologia. 148: 672–681. doi:10.1007/s00442-006-0411-6.

Lima, F.P., Wethey, D.S. 2012. Three decades of high-resolution coastal sea surface temperatures reveal more than warming. Nat Commun. 3: 704. doi:10.1038/ncomms1713.

Lindstrom, S.C. 1973. Marine Benthic Algal Communities in the Flat Top Islands Area of Georgia Strait. Univeristy of British Columbia.

Lindstrom, S.C., Lemay, M.A., Starko, S., Hind, K.R., Martone, P.T. 2021. New and interesting seaweed records from the Hakai area of the central coast of British Columbia, Canada: Chlorophyta. Bot. Mar. 64: 343–361. doi:10.1515/bot-2021-0038.

Little, C., Trowbridge, C.D., Pilling, G.M., Stirling, P., Morritt, D., Williams, G.A. 2017. Long-term fluctuations in intertidal communities in an Irish sea-lough: Limpet-fucoid cycles. ESCA. 196: 70–82. doi:10.1016/j.ecss.2017.06.036.

Mann, K.H. 1973. Seaweeds: Their productivity and strategy for growth. Science. 182: 975–981.

Maxell, B.A., Miller, K.A. 1996. Demographic studies of the annual kelps *Nereocystis luetkeana* and *Costaria costata* (Laminariales, Phaeophyta) in Puget Sound, Washington. Bot. Mar. 39: 479–490. doi:10.1515/botm.1996.39.1-6.479.

McConnico, L.A., Foster, M.S. 2005. Population biology of the intertidal kelp, *Alaria marginata* Postels and Ruprecht: A non-fugitive annual. J. Exp. Mar. Bio. Ecol. 324: 61–75. doi:10.1016/j.jembe.2005.04.006.

Miller, A., Ambrose, R. 2000. Sampling patchy distributions:comparison of sampling designs in rocky intertidal habitats. Mar. Ecol. Prog. Ser. 196: 1–14. doi:10.3354/meps196001.

Milligan, K.L.D., DeWreede, R.E. 2000. Variations in holdfast attachment mechanics with developmental stage, substratum-type, season, and wave-exposure for the intertidal kelp species *Hedophyllum sessile* (C. Agardh) Setchell. J. Exp. Mar. Bio. Ecol. 254: 189–209. doi:10.1016/S0022-0981(00)00279-3.

Mineur, F., Arenas, F., Assis, J., Davies, A.J., Engelen, A.H., Fernandes, F., Malta, E., Thibaut, T., Van Nguyen, T., Vaz-Pinto, F., Vranken, S., Serrão, E.A., De Clerck, O. 2015. European seaweeds under pressure: Consequences for communities and ecosystem functioning. Journal of Sea Research, Protecting Marine Biodiversity to Preserve Ecosystem Functioning: a Tribute to Carlo Heip. J. Sea Res. 98: 91–108.doi:10.1016/j.seares.2014.11.004.

Moy, F.E., Christie, H. 2012. Large-scale shift from sugar kelp (*Saccharina latissima*) to ephemeral algae along the south and west coast of Norway. Mar. Biol. Res. 8: 309–321. doi:10.1080/17451000.2011.637561.

Neilson-Welsh, L. 1999. Saline water intrusion from the Fraser River Estuary: A hydrogeological investigation using field chemical data and a density-dependent groundwater flow model. University of British Columbia.

Nelson, T.A., Haberlin, K., Nelson, A.V., Ribarich, H., Hotchkiss, R., Alstyne, K.L.V., Buckingham, L., Simunds, D.J., Fredrickson, K. 2008. Ecological and physiological controls of species composition in green macroalgal blooms. Ecology. 89: 1287–1298. doi:10.1890/07-0494.1.

Nelson, T.A., Nelson, A.V., Tjoelker, M. 2003. Seasonal and spatial patterns of “Green Tides” (Ulvoid Algal Blooms) and related water quality parameters in the coastal waters of Washington State, USA. 46: 263–275. doi:10.1515/BOT.2003.024.

Nielsen, M.M., Krause-Jensen, D., Olesen, B., Thinggaard, R., Christensen, P.B., Bruhn, A. 2014. growth dynamics of *Saccharina latissima* (Laminariales, Phaeophyceae) in Aarhus Bay, Denmark, and along the species’ distribution range. Mar Biol. 161: 2011–2022. doi:10.1007/s00227-014-2482-y.

Okamoto, D.K., Stekoll, M.S., Eckert, G.L. 2013. Coexistence despite recruitment inhibition of kelps by subtidal algal crusts. 493: 103–112. doi:10.3354/meps10505.

Olabarria, C., Arenas, F., Viejo, R.M., Gestoso, I., Vaz-Pinto, F., Incera, M., Rubal, M., Cacabelos, E., Veiga, P., Sobrino, C. 2013. Response of macroalgal assemblages from rockpools to climate change: effects of persistent increase in temperature and CO2. Oikos. 122: 1065– 1079. doi:10.1111/j.1600-0706.2012.20825.x.

Parmesan, C., Yohe, G. 2003. A globally coherent fingerprint of climate change impacts across natural systems. Nature. 421: 37–42. doi:10.1038/nature01286.

Pawluk, K.A. 2016. Impacts and interactions of two non-indigenous seaweeds *Mazzaella japonica* (Mikami) Hommersand and *Sargassum muticum* (Yendo) Fensholt in Baynes Sound, British Columbia (Thesis).

Pfister, C.A., Altabet, M.A., Weigel, B.L. 2019. Kelp beds and their local effects on seawater chemistry, productivity, and microbial communities. Ecology. 100: e02798. doi:10.1002/ecy.2798.

Poloczanska, E.S., Brown, C.J., Sydeman, W.J., Kiessling, W., Schoeman, D.S., Moore, P.J., Brander, K., Bruno, J.F., Buckley, L.B., Burrows, M.T., Duarte, C.M., Halpern, B.S., Holding, J., Kappel, C.V., O’Connor, M.I., Pandolfi, J.M., Parmesan, C., Schwing, F., Thompson, S.A., Richardson, A.J. 2013. Global imprint of climate change on marine life. Nat. Clim. Change. 3: 919–925. doi:10.1038/nclimate1958.

R Core Team. 2022. R: A Language and Environment for Statistical Computing. R Foundation for Statistical Computing, Vienna, Austria.

Rawal, D.S., Kasel, S., Keatley, M.R., Nitschke, C.R. 2015. Herbarium records identify sensitivity of flowering phenology of eucalypts to climate: Implications for species response to climate change. Austral Ecol. 40: 117–125. doi:10.1111/aec.12183.

Raymond, A.E.T. 2020. Life cycles of the kelps *Saccharina latissima* and *Alaria marginata*: implications for mariculture and ecology in Alaska (Thesis).

Rogers-Bennett, L., Catton, C.A. 2019. Marine heat wave and multiple stressors tip bull kelp forest to sea urchin barrens. Sci Rep. 9: 15050. doi:10.1038/s41598-019-51114-y.

Ryan, S.A., Wohlgeschaffen, G.D., Jahan, N., Niu, H., Ortmann, A.C., Brown, T.N., King, T.L., Clyburne, J. 2019. State of knowledge on fate and behaviour of ship-source petroleum product spills. Fisheries and Oceans Canada = Pêches et océans Canada, Ottawa.

Salvaterra, T., Green, D.S., Crowe, T.P., O’Gorman, E.J. 2013. Impacts of the invasive alga *Sargassum muticum* on ecosystem functioning and food web structure. Biol. Invasions. 15: 2563–2576. doi:10.1007/s10530-013-0473-4.

Saunders, G., Millar, K. 2014. A DNA barcode survey of the red algal genus *Mazzaella* in British Columbia reveals overlooked diversity and new distributional records: Descriptions of *M. dewreedei* sp. nov. and *M. macrocarpa* sp. nov. Botany. 92: doi:10.1139/cjb-2013-0283.

Silva, P., Woodfield, R., Cohen, A., Harris, L., Goddard, J. 2002. First Report of the Asian kelp *Undaria pinnatifida* in the Northeastern Pacific Ocean. Biol. Invasions. 4: 333–338. doi:10.1023/A:1020991726710.

Starko, S., Bailey, L.A., Creviston, E., James, K.A., Warren, A., Brophy, M.K., Danasel, A., Fass, M.P., Townsend, J.A., Neufeld, C.J. 2019. Environmental heterogeneity mediates scale-dependent declines in kelp diversity on intertidal rocky shores. PLoS One. 14: e0213191. doi:10.1371/journal.pone.0213191.

Steneck, R.S., Graham, M.H., Bourque, B.J., Corbett, D., Erlandson, J.M., Estes, J.A., Tegner, M.J. 2002. Kelp forest ecosystems: biodiversity, stability, resilience and future. Envir. Conserv. 29: 436–459. doi:10.1017/S0376892902000322.

Suryan, R.M., Arimitsu, M.L., Coletti, H.A., Hopcroft, R.R., Lindeberg, M.R., Barbeaux, S.J., Batten, S.D., Burt, W.J., Bishop, M.A., Bodkin, J.L., Brenner, R., Campbell, R.W., Cushing, D.A., Danielson, S.L., Dorn, M.W., Drummond, B., Esler, D., Gelatt, T., Hanselman, D.H., Hatch, S.A., Haught, S., Holderied, K., Iken, K., Irons, D.B., Kettle, A.B., Kimmel, D.G., Konar, B., Kuletz, K.J., Laurel, B.J., Maniscalco, J.M., Matkin, C., McKinstry, C.A.E., Monson, D.H., Moran, J.R., Olsen, D., Palsson, W.A., Pegau, W.S., Piatt, J.F., Rogers, L.A., Rojek, N.A., Schaefer, A., Spies, I.B., Straley, J.M., Strom, S.L., Sweeney, K.L., Szymkowiak, M., Weitzman, B.P., Yasumiishi, E.M., Zador, S.G. 2021. Ecosystem response persists after a prolonged marine heatwave. Sci Rep. 11: 6235. doi:10.1038/s41598-021-83818-5.

Teagle, H., Hawkins, S.J., Moore, P.J., Smale, D.A. 2017. The role of kelp species as biogenic habitat formers in coastal marine ecosystems. J. Exp. Mar. Biol. Ecol. 492: 81–98. doi:10.1016/j.jembe.2017.01.017.

Thom, R.M., Albright, R.G. 1990. Dynamics of benthic vegetation standing-stock, irradiance, and water properties in central Puget Sound. Mar. Biol. 104: 129–141. doi:10.1007/BF01313166.

Todd, P.A., Heery, E.C., Loke, L.H.L., Thurstan, R.H., Kotze, D.J., Swan, C. 2019. Towards an urban marine ecology: characterizing the drivers, patterns and processes of marine ecosystems in coastal cities. Oikos. 128: 1215–1242. doi:10.1111/oik.05946.

Tsleil-Waututh Nation, 2017. Burrard Inlet Action Plan (2017 version).

Twist, B.A., Mazel, F., Zaklan Duff, S., Lemay, M.A., Pearce, C.M., Martone, P.T. 2024. Kelp and sea urchin settlement mediated by biotic interactions with benthic coralline algal species. J Phyc. 60: 363–379. doi:10.1111/jpy.13420.

Ulaski, B.P., Konar, B., Otis, E.O. 2020. Seaweed reproduction and harvest rebound in Southcentral Alaska: Implications for wild stock management. ESCO. 43: 2046–2062. doi:10.1007/s12237-020-00740-1.

Van Alstyne, K.L. 2016. Seasonal changes in nutrient limitation and nitrate sources in the green macroalga *Ulva lactuca* at sites with and without green tides in a northeastern Pacific embayment. Marine Pollution Bulletin. 103: 186–194. doi:10.1016/j.marpolbul.2015.12.020.

Vergés, A., Campbell, A.H., Wood, G., Kajlich, L., Eger, A.M., Cruz, D., Langley, M., Bolton, D., Coleman, M.A., Turpin, J., Crawford, M., Coombes, N., Camilleri, A., Steinberg, P.D., Marzinelli, E.M. 2020. Operation Crayweed: Ecological and sociocultural aspects of restoring Sydney’s underwater forests. Ecol. Manag. Restor. 21: 74–85. doi:10.1111/emr.12413.

Weitzman, B., Konar, B., Iken, K., Coletti, H., Monson, D., Suryan, R., Dean, T., Hondolero, D., Lindeberg, M. 2021. Changes in Rocky Intertidal Community Structure During a Marine Heatwave in the Northern Gulf of Alaska. Front. Mar. Sci. 8: 2296–7745. doi: 10.3389/fmars.2021.556820.

Wernberg, T., Smale, D.A., Thomsen, M.S. 2012. A decade of climate change experiments on marine organisms: procedures, patterns and problems. Glob. Chang. Biol. 18: 1491–1498. doi:10.1111/j.1365-2486.2012.02656.x.

Wernberg, T., Thomsen, M.S., Tuya, F., Kendrick, G.A., Staehr, P.A., Toohey, B.D. 2010. Decreasing resilience of kelp beds along a latitudinal temperature gradient: potential implications for a warmer future. Ecol. Lett. 13: 685–694. doi:10.1111/j.1461-0248.2010.01466.x.

Whalen, M.A., Starko, S., Lindstrom, S.C., Martone, P.T. 2023. Heatwave restructures marine intertidal communities across a stress gradient. Ecology. 104: e4027. doi:10.1002/ecy.4027.

White, R.H., Anderson, S., Booth, J.F., Braich, G., Draeger, C., Fei, C., Harley, C.D.G., Henderson, S.B., Jakob, M., Lau, C.-A., Mareshet Admasu, L., Narinesingh, V., Rodell, C., Roocroft, E., Weinberger, K.R., West, G. 2023. The unprecedented Pacific Northwest heatwave of June 2021. Nat. Com. 14: 727. doi:10.1038/s41467-023-36289-3.

Wickham, H. 2016. ggplot2: Elegant Graphics for Data Analysis. Springer-Verlag New York. Wickham, H. 2011. The Split-Apply-Combine Strategy for Data Analysis. J. Stat. Softw. 40: 1–29.

Wonham, M., Gerstle, C., Bates, C. 2023. Combining current and historical biodiversity surveys reveals order of magnitude greater richness in a British Columbia marine protected area. Can. Field. Nat. 136: 348–360. doi:10.22621/cfn.v136i4.2903.

Zabin, C., Ashton, G., Brown, C., Ruiz, G. 2009. Northern range expansion of the Asian kelp *Undaria pinnatifida* (Harvey) Suringar (Laminariales, Phaeophyceae) in western North America. AI. 4: 429–434. doi:10.3391/ai.2009.4.3.1.

